# Benchmarking Machine Learning Methods for Antimicrobial Peptide Prediction and Discovery in Ancient Microbiomes

**DOI:** 10.64898/2026.11.17.767431

**Authors:** Quan Zhao, Qin Ma, Jie Luo, Can Chen, Shanlin Ke

## Abstract

Antimicrobial peptides (AMPs) offer a promising alternative to conventional antibiotics amid the growing global threat of antimicrobial resistance. Expanding AMP discovery requires both effective computational tools to accurately identify candidate peptides and access to underexplored biological resources that may harbor novel AMPs. However, the comparative performance of existing computational methods remains poorly understood, while ancient microbiomes have received limited attention as potential reservoirs of AMPs. Here, we systematically benchmarked six machine learning methods across with twelve feature configurations spanning sequence, structural, and protein language model features under a unified framework using eight curated AMP datasets. The best-performing methods were subsequently applied to three ancient metagenomic datasets representing the human gut (archaeological fecal sediments), human oral cavity (dental calculus), and mammoth-associated communities (teeth, skulls, and skin). In parallel, we employed reinforcement learning to select candidate peptides directly from ancient sequence pools without relying on modern AMP labels. Our benchmarking shows that protein language model embeddings provide the most informative features, and deep learning models, including convolutional, transformer-based, and recurrent architectures, outperform classical methods. Applying these models to ancient microbiomes identified candidates occupied the physicochemical space characteristic of AMPs but differed from those of candidates predicted from modern microbiome catalogs. We identified 111, 76, and 50 high-confidence candidate AMPs from the Neanderthal gut, ancient oral, and mammoth-associated communities, respectively. Reinforcement learning selected additional candidate AMPs with distinct properties compared with those predicted by models trained on modern AMP databases. Together, these findings establish a systematic benchmark for AMP prediction and highlight ancient microbiomes as an untapped source of antimicrobial peptide candidates. By integrating supervised machine learning with reinforcement learning, this study expands AMP discovery beyond modern microbiome catalogs and provides new opportunities to identify potentially novel AMPs.

## Introduction

Antimicrobial resistance (AMR) has become one of the most serious threats to global public health, with approximately 1.27 million deaths attributed directly to bacterial AMR in 2019^1^ and projections reaching as many as 10 million annually by 2050, surpassing cancer as a leading cause of death worldwide^2^. However, the development of new antibiotics has remained persistently slow, leaving a widening gap between clinical need and available therapeutics^1^. Antimicrobial peptides (AMPs) have attracted growing interest as a promising alternative for combating AMR. In contrast to conventional antibiotics, which typically inhibit a single molecular target, AMPs act through diverse and complementary mechanisms, including direct disruption of microbial membranes, translocation into cells to interfere with essential intracellular processes, and recruitment and modulation of host immune responses^3, 4^. These diverse mechanisms, together with immunomodulatory activity, can confer broad-spectrum antimicrobial effects and reduce the propensity for resistance development^5^. Discovering new AMPs at scale depends on two main factors: the computational tools used to identify candidate AMPs and the biological sources for which they are mined.

A wide range of computational tools has been developed to distinguish AMPs from non-antimicrobial sequences, spanning homology-based searches, classical machine learning classifiers, and, more recently, deep learning and protein language model methods^6–11^. Early approaches relied on sequence similarity and motif matching to known AMPs, as in AntiBP2^12^. Classical machine learning classifiers combined handcrafted sequence and physicochemical descriptors with algorithms such as support vector machines and random forests, as in CAMP^13^ and AmPEP^7^. More recent deep learning and protein language model methods learned representations directly from sequence, including the convolutional-recurrent AMP Scanner^8^, the attention-based AMPlify^10^, and transformer-based models such as AMP-BERT^14^. Several studies have compared such predictors: Gabere and Noble benchmarked ten web-based tools on common test sets^15^, Xu et al. assessed 34 methods using an independent test set and six validation datasets^6^, and Sidorczuk et al. showed that such benchmarks can be biased by how non-AMP negative sequences are selected^16^, with more recent efforts evaluating predictors under standardized frameworks^17^. These studies have largely compared existing tools as integrated end-to-end predictors, rather than disentangling the contributions of the underlying feature representations and model architectures under a common training and evaluation protocol. It therefore remains unclear which feature representations and which model architectures most reliably distinguish AMPs from non-antimicrobial sequences, particularly given the rapid proliferation of feature encodings and modern architectures, including protein language models, whose relative strengths have not been established on common ground.

The second factor, the choice of source, has received comparatively little attention: computational AMP mining has to date focused overwhelmingly on modern microbiomes^18–21^. Yet contemporary microbial communities have experienced substantial losses of diversity under industrialized diets, hygiene practices, and pervasive antibiotic exposure, and may no longer harbor antimicrobial molecules that were once widespread^22^. Ancient microbiomes offer a compelling and largely untapped alternative. Preserved in substrates such as coprolites, paleofeces, and dental calculus, ancient metagenomes capture microbial communities that existed before the widespread clinical and agricultural use of antibiotics^23, 24^, and may consequently provide access to antimicrobial molecules associated with microbial competition in ecological environments predating modern antibiotic selection^22, 25^. A recent study has begun to apply machine learning to ancient communities^26^, but examined a single source of palaeofaeces, leaving the broader diversity of preserved microbial communities largely unexplored. Importantly, supervised predictors are trained on modern labeled datasets and may therefore favor sequence patterns represented in those datasets, potentially overlooking ancient peptides with divergent sequence characteristics. This limitation motivates complementary approaches that can explore ancient sequence pools without relying on modern AMP labels.

To address these gaps, we first established a unified benchmark that systematically compares a broad set of feature representations and state-of-the-art prediction methods for AMP identification across eight curated datasets, providing a systematic assessment of their ability to distinguish AMPs from non-antimicrobial sequences. We then applied the best-performing methods with the most informative single feature representation identified by the benchmark, to three ancient metagenomes: a Neanderthal gut microbiome recovered from fecal sediments, an ancient oral microbiome recovered from dental calculus, and a mammoth-associated microbiome recovered from skeletal and dental remains. In parallel, we explored a complementary approach by leveraging reinforcement learning to select candidates from the same ancient sequence pools based on reward functions associated with physicochemical criteria alone, without using modern AMP labels during the search. Through these complementary approaches, we identified high confidence AMP candidates from these ancient communities and characterized their physicochemical properties. By pairing a rigorous prediction benchmark with the exploration of ancient microbial sequence space, this work provides both a reusable foundation for AMP prediction and a route to antimicrobial candidates that lie outside the scope of present-day sequence collections, offering new starting points for combating AMR.

## Results

### A unified framework for AMP prediction

To systematically evaluate the performance of different computational approaches for identifying AMPs from biological sequence, we developed a unified framework that pairs a diverse set of sequence representations with a diverse set of prediction methods under a consistent benchmarking conditions (**Figure 1**). The framework consists of two components: numerical representations of peptide sequences and classification methods for predicting antimicrobial activity.

**Figure 1.**
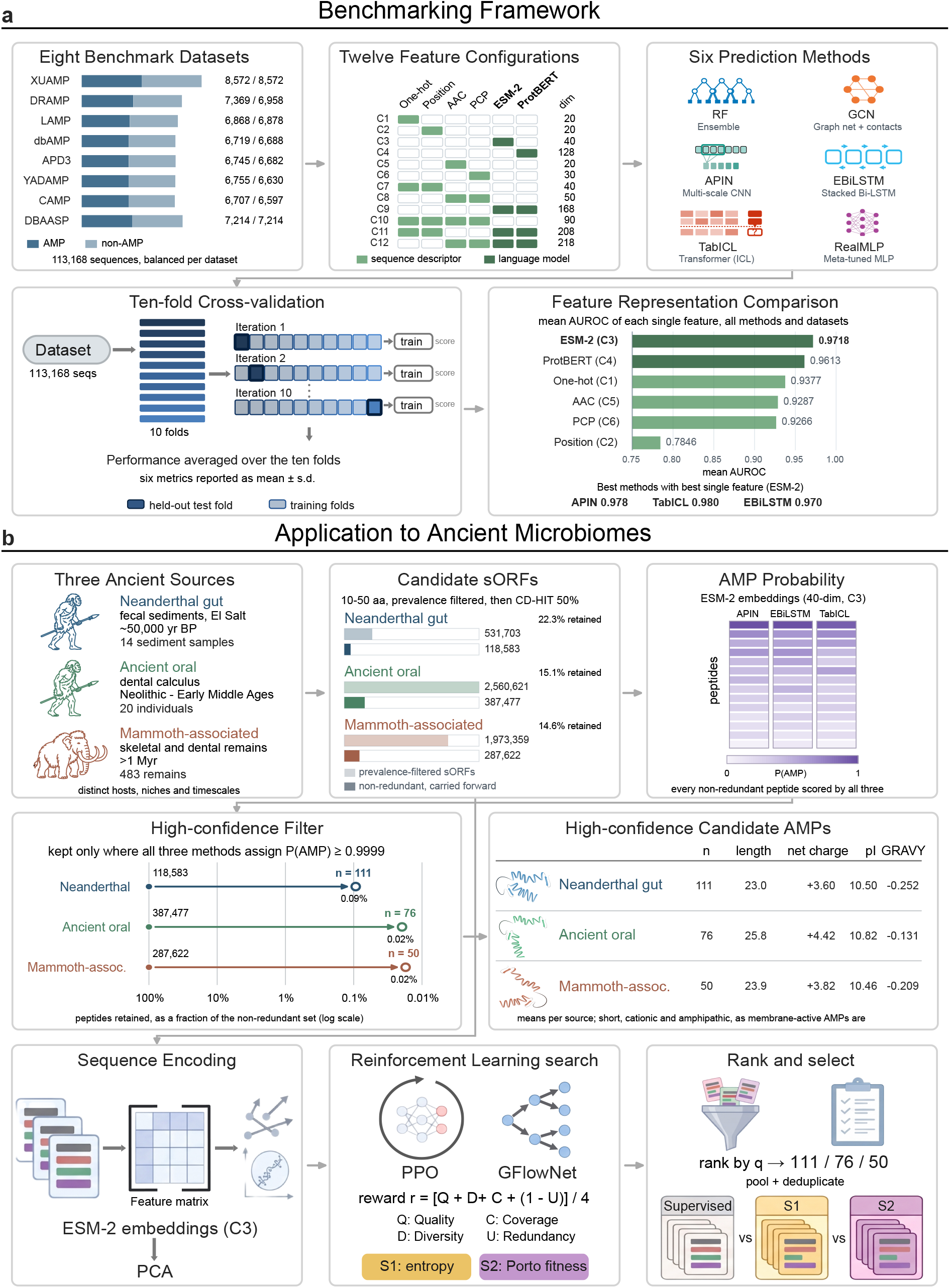
Overview of the benchmarking framework and its application to ancient microbiomes. **(a)** Benchmarking framework. Six machine learning methods were evaluated across eight curated AMP benchmark datasets and twelve feature configurations. Reading left to right and then top to bottom: the eight datasets, each with its balanced counts of AMP and non-AMP sequences; the twelve feature configurations of Table 1, formed from six feature types (one-hot encoding, sinusoidal position encoding, amino acid composition, physicochemical properties, and ESM-2 and ProtBERT language model embeddings) used individually and in combination, with the dimensionality of each configuration; the six prediction methods (RF, GCN, APIN, EBiLSTM, TabICL, and RealMLP); ten-fold cross-validation used for evaluation; and a comparison of the six single-feature representations by mean AUROC, in which ESM-2 was the most informative single feature. C3, ESM-2 embeddings alone, provided the feature representation carried forward to the ancient application. The three methods carried forward (APIN, TabICL, and EBiLSTM) are listed with their mean AUROC at C3. **(b)** Application to ancient microbiomes. The benchmarked pipeline was applied to three ancient metagenomic datasets, each identified by color throughout. Reading left to right and then top to bottom: the three ancient sources, a Neanderthal gut microbiome, an ancient human oral microbiome, and a mammoth-associated microbiome, with the number of samples analyzed in each source study; prevalence filtering and clustering of candidate short open reading frames (sORFs) at 50% sequence identity, with the number of sequences before and after clustering. From these non-redundant sORF pools the analysis proceeds along two routes. In the supervised route, each peptide is scored with the three selected methods on ESM-2 embeddings to obtain per-peptide AMP probabilities, and a high-confidence filter retains peptides for which all three methods assigned a probability of at least 0.9999, shown as the fraction of each non-redundant set that survives, yielding high-confidence candidate AMPs numbering 111, 76, and 50 for the Neanderthal gut, ancient oral, and mammoth-associated microbiomes, respectively, with their mean length, net charge, isoelectric point and GRAVY. In the reinforcement-learning route, the same sORF pools are encoded (sequence descriptors reduced by PCA and L2-normalized) and searched with PPO and GFlowNet under two reward designs (S1, composition entropy; S2, physicochemical fitness), without modern AMP labels, and the visited sequences are ranked and selected to give candidate sets compared against the supervised candidates. The receiver operating characteristic curves, the fold allocation and the per-peptide probability heat map are schematic and are drawn for illustration; all counts, dimensionalities and summary statistics shown are the values reported in the text.

**Table 1:**
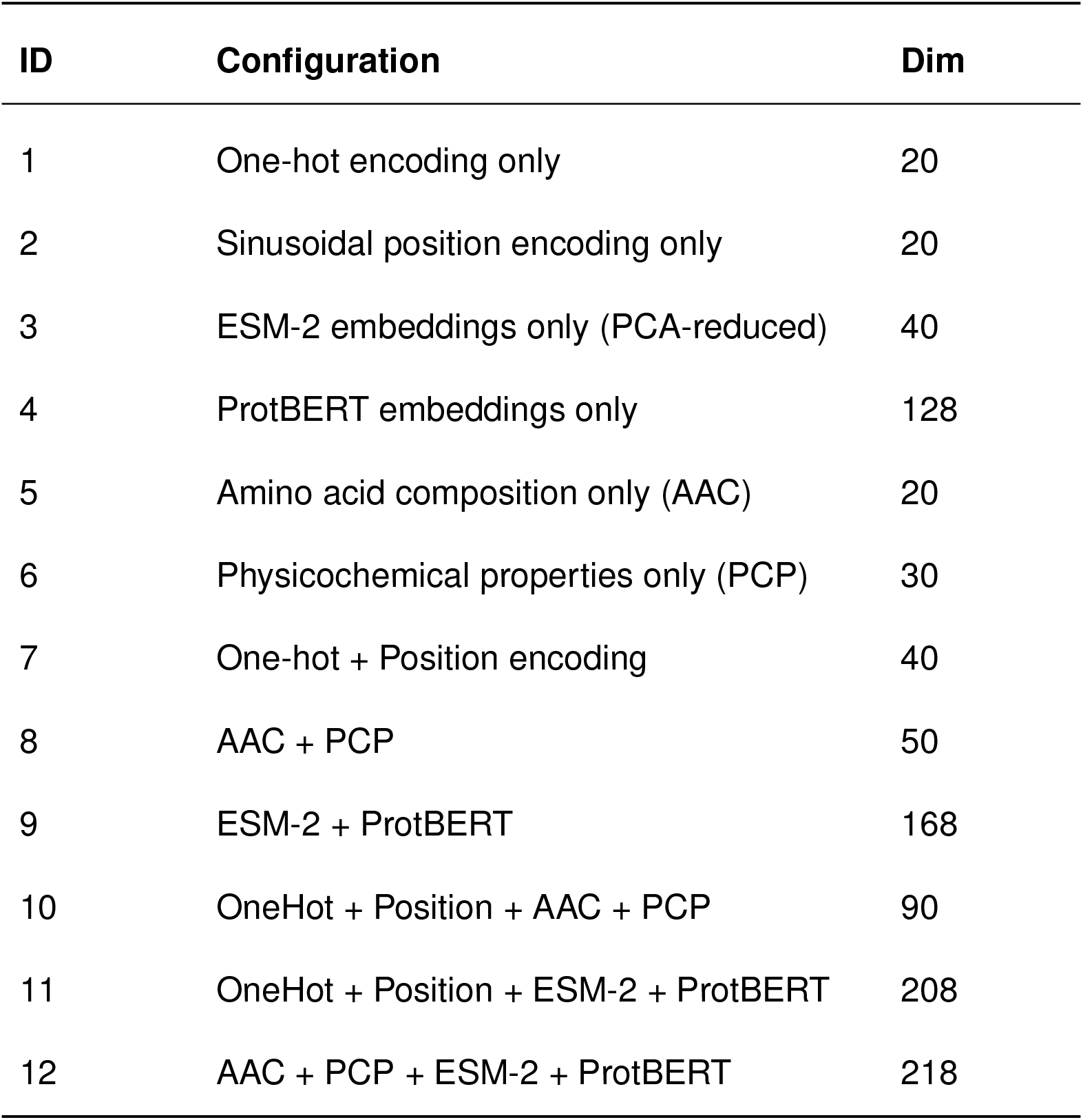
Summary of feature configurations.

We compared three complementary families for peptide representations. The first is handcrafted descriptors, which summarize a peptide using properties long known to matter for antimicrobial activity, its amino acid composition, physicochemical characteristics such as charge and hydrophobicity, and the identity and position of each residue along the sequence. The second is structural information through predicted contacts between residues that approximate how the peptide folds. The third is protein language model embeddings learned from large-scale protein sequence datasets, which can capture evolutionary and contextual information beyond predefined descriptors. We evaluated these representations individually and in representative combinations, yielding twelve feature configurations (Table 1).

For prediction, we selected six methods that span the major paradigms used for sequence classification: a tree-based ensemble, a graph neural network that operates on predicted residue contacts, a convolutional network that scans the sequence for informative motifs, a bidirectional recurrent network that reads the sequence in both directions, a meta-tuned multilayer perceptron, and a tabular in-context transformer that learns directly from labeled examples without conventional model training (Table 2). These methods span traditional machine learning and modern deep learning, including both conventional model-training and in-context learning approaches. Evaluating all six methods against all twelve representations across eight curated AMP datasets provides a comprehensive picture of which strategies are most effective. Detailed mathematical descriptions of representations and methods are provided in the Methods and Supplementary Note 1.

**Table 2:** Summary of prediction methods.

| **Method** | **Type** | **Category** | **Structure** |
|---|---|---|---|
| Random Forest | Ensemble | Traditional ML | No |
| GCN | Graph Neural Network | Deep Learning | Yes |
| APIN | Multi-scale CNN | Deep Learning | No |
| EBiLSTM | Stacked Bi-LSTM | Deep Learning | No |
| TabICL | Transformer (ICL) | Foundation Model | No |
| RealMLP | Meta-tuned MLP | Deep Learning | No |

### Protein language model embeddings are the most informative features for AMP prediction

We first evaluated the ability of different feature representations to distinguish AMPs from non-antimicrobial sequences, using the area under the receiver operating characteristic curve (AUC) as the primary performance meteric. As shown in **Figure 2**, among the six individual feature types, ESM-2 embeddings^11^ achieved the highest average AUC of 0.9718 across all methods and datasets, followed by ProtBERT embeddings^27^ (0.9613), one-hot encoding (0.9377), amino acid composition (0.9287), physicochemical properties (0.9266), and sinusoidal position encoding^28^ (0.7846). The two protein language model embeddings (ESM-2 and ProtBERT) outperformed the handcrafted features (one-hot, AAC, and PCP) by a wide margin, potentially reflecting their ability to capture the evolutionary and contextual information learned from large protein sequence databases. Sinusoidal positional encoding alone performed far worse than every other feature type, indicating that positional information along provides limited discriminative information for AMP classification.

**Figure 2.**
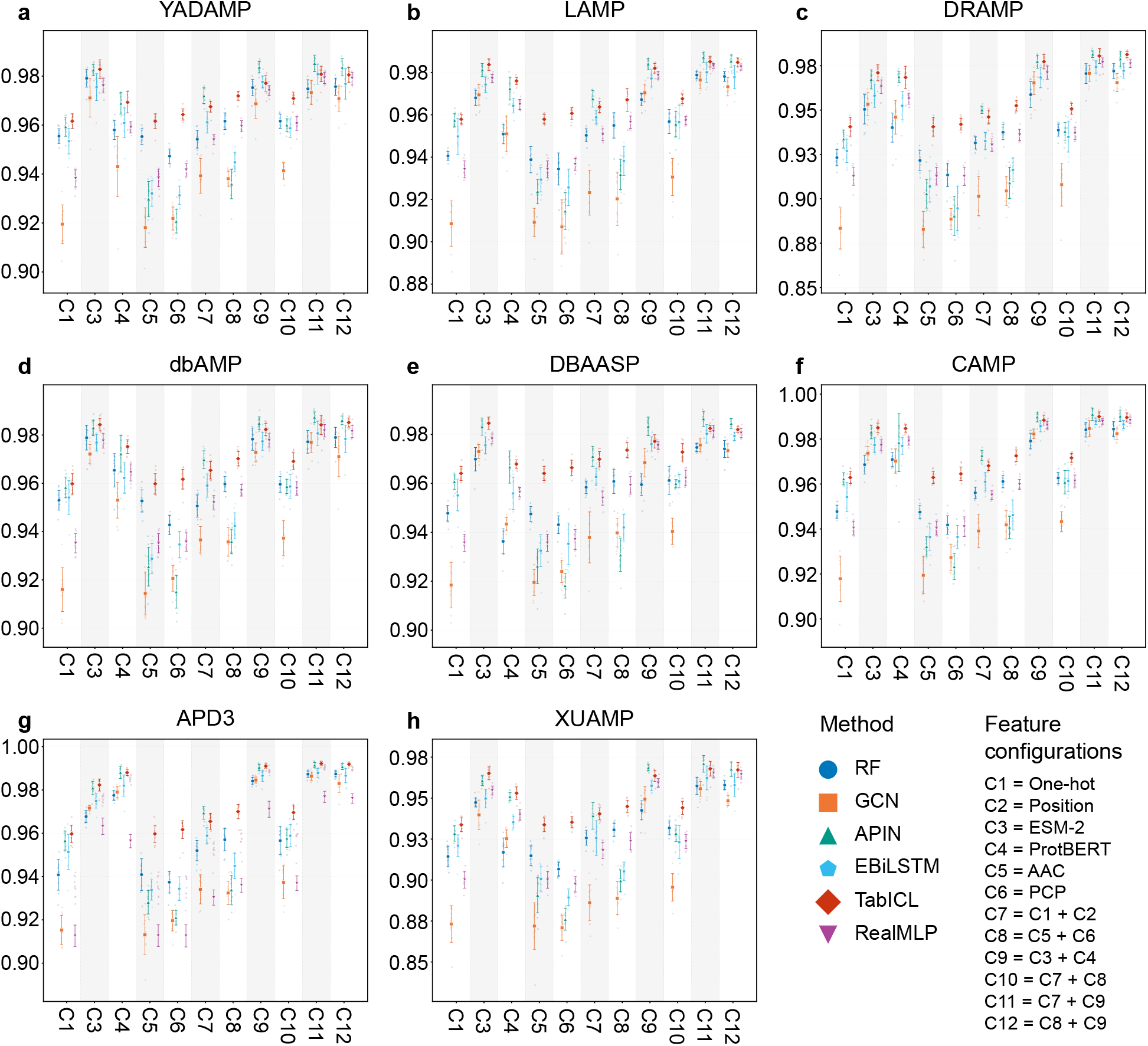
AUROC performance comparison of six machine learning methods across eight AMP benchmark datasets. Each panel (**a**–**h**) corresponds to one benchmark dataset and displays the area under the receiver operating characteristic curve (AUROC) for RF (Random Forest), GCN (Graph Convolutional Network), APIN (CNN-based), EBiLSTM (Enhanced Bidirectional LSTM), TabICL (Transformer-based), and RealMLP across 11 feature configurations. Configurations are labelled C1–C12 as defined in Table 1; the position encoding (C2) is omitted here for clarity. Error bars represent standard deviation from 10-fold cross-validation, with individual fold results shown as lighter background points.

Combining features generally improved performance over individual features (**Figure 2**). The two best-performing configurations were Configuration 11 (One-hot + Position + ESM-2 + ProtBERT, 208 dimensions, average AUC = 0.9795) and Configuration 12 (ESM-2 + ProtBERT + PCP + AAC, 218 dimensions, average AUC = 0.9781). Both configurations incorporate the two protein language model embeddings, indicating that ESM-2 and ProtBERT capture complementary information. Interestingly, the addition of traditional features (one-hot and position encoding in Configuration 11, or PCP and AAC in Configuration 12) to the language model embeddings provided only modest improvements over ESM-2 + ProtBERT alone (Configuration 9, average AUC = 0.9752), suggesting that the language model embeddings already encode much of the predictive information represented by handcrafted descriptors. Among the traditional feature combinations, Configuration 10 (One-hot + Position + PCP + AAC, 90 dimensions, average AUC = 0.9502) and Configuration 8 (PCP + AAC, 50 dimensions, average AUC = 0.9423) showed improved performance compared with individual traditional features but remained substantially below configurations that included protein language model embeddings. This gap persisted across all methods and datasets, reinforcing the finding that protein language model features are the most informative feature representations evaluated in this benchmark.

### APIN achieves the best overall performance across methods and datasets

We next compared six prediction methods (Table 2) chosen to represent the major paradigms applied to biological sequence classification: a tree-based ensemble (Random Forest^29^), a graph neural network operating on predicted residue contacts (GCN^30^), a multi-scale convolutional network (APIN^9^), a bidirectional recurrent network (EBiLSTM^31^), a meta-tuned multilayer perceptron (RealMLP^32^), and a tabular in-context transformer (TabICL^33^). This selection spans tabular and sequence-aware architectures, as well as conventional training and in-context learning paradigms, allowing us to assess how architectural inductive biases interact with the feature representations described above.

When evaluated on the best-performing feature configuration (Configuration 11), the six methods ranked as follows by average AUC across all eight datasets: APIN (0.9851), TabICL (0.9829), EBiLSTM (0.9793), RealMLP (0.9790), Random Forest (0.9756), and GCN (0.9750). APIN achieved the highest AUC on seven of the eight datasets, while TabICL achieved the highest AUC on APD3 (0.9922 vs. APIN’s 0.9915). The per-dataset AUC of each method across all feature configurations is presented in **Figure 2**. To test whether these differences were statistically meaningful, we applied a Friedman test to the per-dataset AUCs of the six methods at Configuration 11, followed by a Nemenyi post-hoc test. The methods differed significantly overall (Friedman *χ*^2^ = 34.79, df = 5, *p* = 1.7 *×* 10*^−^*^6^). By average rank, APIN was the top performer (mean rank 1.12), followed by TabICL (1.94), EBiLSTM (3.56), RealMLP (3.62), RF (5.25), and GCN (5.50). The Nemenyi test yielded a critical difference of 2.67 (*α* = 0.05), meaning that methods whose mean ranks differ by more than this value are significantly different. APIN and TabICL thus ranked significantly above Random Forest and GCN, whereas the four top-ranked methods (APIN, TabICL, EBiLSTM, and RealMLP), with mean ranks spanning 1.12 to 3.62, fell within one critical difference of each other and did not differ significantly.

The relative performance of the methods reflects their architectural inductive biases. APIN’s multi-scale convolutional design, with filter sizes from 2 to 28 residues, is well suited to capturing sequence motifs of varying lengths, which likely underlies its top performance. TabICL ranked second despite operating without gradient-based fine-tuning, relying instead on in-context learning from labeled examples in its context window. EBiLSTM followed closely, preserving positional information by processing the full residue sequence through bidirectional layers rather than pooling residues into a single vector, and it consistently matched or exceeded the tree-based and graph-based methods. RealMLP performed comparably, transferring the effective default hyperparameters from its meta-learning optimization to AMP prediction without per-task tuning. Notably, Random Forest performed competitively despite being the simplest method and remained stable across feature configurations, indicating that informative feature representations can limit the gains from additional architectural complexity. GCN, the only method to use predicted residue contact maps, ranked lowest, which may reflect that the contact maps add little beyond the residue-level features already captured, or that contact prediction is unreliable for short peptides and confers no structural advantage. Overall, the strongest results came from methods that model the peptide sequence directly, whereas explicit structural modeling offered no clear benefit at this sequence length.

### Method rankings are consistent across datasets of differing size and composition

Predictive performance varied across datasets, but this variation tracked how each dataset was assembled, whether aggregated from multiple sources or individually curated, rather than its size. XUAMP, which aggregates sequences from several databases (DRAMP, LAMP, and YADAMP) with redundancy removal, was consistently the most challenging dataset: it gave the lowest AUC for every method. For instance, APIN achieved an AUC of 0.9709 with Configuration 11, compared with 0.9915 on APD3 and 0.9910 on CAMP. The individually curated databases generally showed higher AUCs, and their performance did not track dataset size; APD3 and CAMP, which achieved the highest scores, are similar in size to the other single source datasets and not markedly smaller than XUAMP. This pattern likely reflects the greater sequence diversity and reduced redundancy of the aggregated XUAMP benchmark, which may make classification more challenging, rather than any simple effect of sample count. Despite these differences in absolute performance, the relative ranking of methods remained largely consistent across all eight datasets, with APIN and TabICL occupying the top two positions in most cases. This consistency suggests that the relative performance advantages of these methods were not limited to a particular dataset, lending confidence to the robustness of the benchmarking results across datasets with different characteristics.

### AMP discovery in ancient microbiomes using models trained on modern data

Having established the relative performance of the prediction methods and feature representations, we next applied the benchmarked framework to explore the potential of ancient microbiomes as reservoirs of candidate AMPs. We analyzed three ancient metagenomic datasets: 1) the Neanderthal gut microbiome reconstructed from Middle Paleolithic fecal sediments dated to approximately 50,000 years ago (n = 14 sediment samples)^34^; 2) an ancient human oral microbiome preserved in archaeological dental calculus spanning the Neolithic to the Early Middle Ages (n = 20 individuals)^35^; and 3) a mammoth-associated microbiome recovered from skeletal and dental remains spanning more than one million years (n = 483 remains)^36^.

Following quality control and removal of host-derived sequences, we assembled metagenomic reads into longer contigs and identified small open reading frames (sORFs) encoding peptides 10–50 amino acids in length. To prioritize reproducibly detected candidates, we retained sORF sequences detected in at least 10% of samples within a dataset or at least 5 independent samples, whichever criterion was less stringent, yielding 531,703, 2,560,621, and 1,973,359 sORFs, respectively. Clustering at 50% sequence identity reduced these to 118,583 sequences for the Neanderthal gut, 387,477 for the ancient oral microbiome, and 287,622 for the mammoth-associated dataset. We next assessed sequence overlap with matched modern references: HRGM2 for Neanderthal gut^37^, HROM for ancient oral^38^, and Asian elephant gut microbiome for the mammoth-associated dataset^39^. Only 0.21% of Neanderthal gut and 0.03% of mammoth-associated sORFs had close modern counterparts, compared with 58.1% of ancient oral sORFs (Supplementary Note 2). Thus, ancient gut and mammoth-associated sORFs were largely distinct from their matched modern references, whereas most ancient oral sORFs had modern counterparts.

We then scored non-redundant peptides with the three top-performing methods from our benchmark (APIN, TabICL, and EBiLSTM), using ESM-2 embeddings alone as the feature representation. Although the full multi-feature combination achieved marginally higher benchmark performance, ESM-2 was the most informative individual feature representation and provided near-equivalent predictive performance while substantially reducing the computational cost of scoring hundreds of thousands of candidate peptides. For the ancient application, each method was trained on a stratified 90/10 split of the combined eight-dataset training set, with the 10% partition held out for early stopping rather than used in cross-validation. Performance on this held-out partition was consistent with the benchmark results, with AUCs of 0.9661 for APIN, 0.9654 for TabICL, and 0.9653 for EBiLSTM. AMP prediction rates for APIN, TabICL, and EBiLSTM were 48.2%, 59.3%, and 88.9% in Neanderthal gut (43.4%; 51,455 peptides unanimously predicted), 40.7%, 54.2%, and 85.7% in ancient oral (36.2%; 140,155 peptides), and 35.8%, 52.4%, and 87.3% in mammoth-associated samples (30.8%; 88,690 peptides). Notably, at every mean probability threshold of 0.90 or above, the unanimous set and the full set were identical, indicating that any peptide scoring highly on average was already called by all three methods.

We next compared the three-model consensus predictions with AMPSphere, a catalogue of 863,498 AMPs predicted from modern global microbiomes^20^, at both the sequence and physicochemical levels. At the sequence level, fewer than 0.03% of the ancient consensus predictions had a close match in AMPSphere (Supplementary Figure 2), indicating they are largely novel relative to modern predicted AMPs. Physicochemically, they fell within the same overall space as the catalogue (**Figure 3**): short, cationic, and amphipathic. Relative to AMPSphere, however, they were on average slightly shorter and modestly less cationic, and notably more hydrophilic, with mean GRAVY values near *−*0.3 compared with approximately +0.05 for the modern catalogue. Given the very large sample sizes, nearly all pairwise differences were statistically significant (two-sided Mann-Whitney *U*), so we interpret the comparison through the location and spread of each distribution rather than through significance. Thus, although the ancient sORFs are largely distinct in sequence from modern sORFs, the peptides nominated as AMPs share the core physicochemical characteristics of modern AMPs, indicating a novel repertoire with conserved biochemical hallmarks.

**Figure 3.**
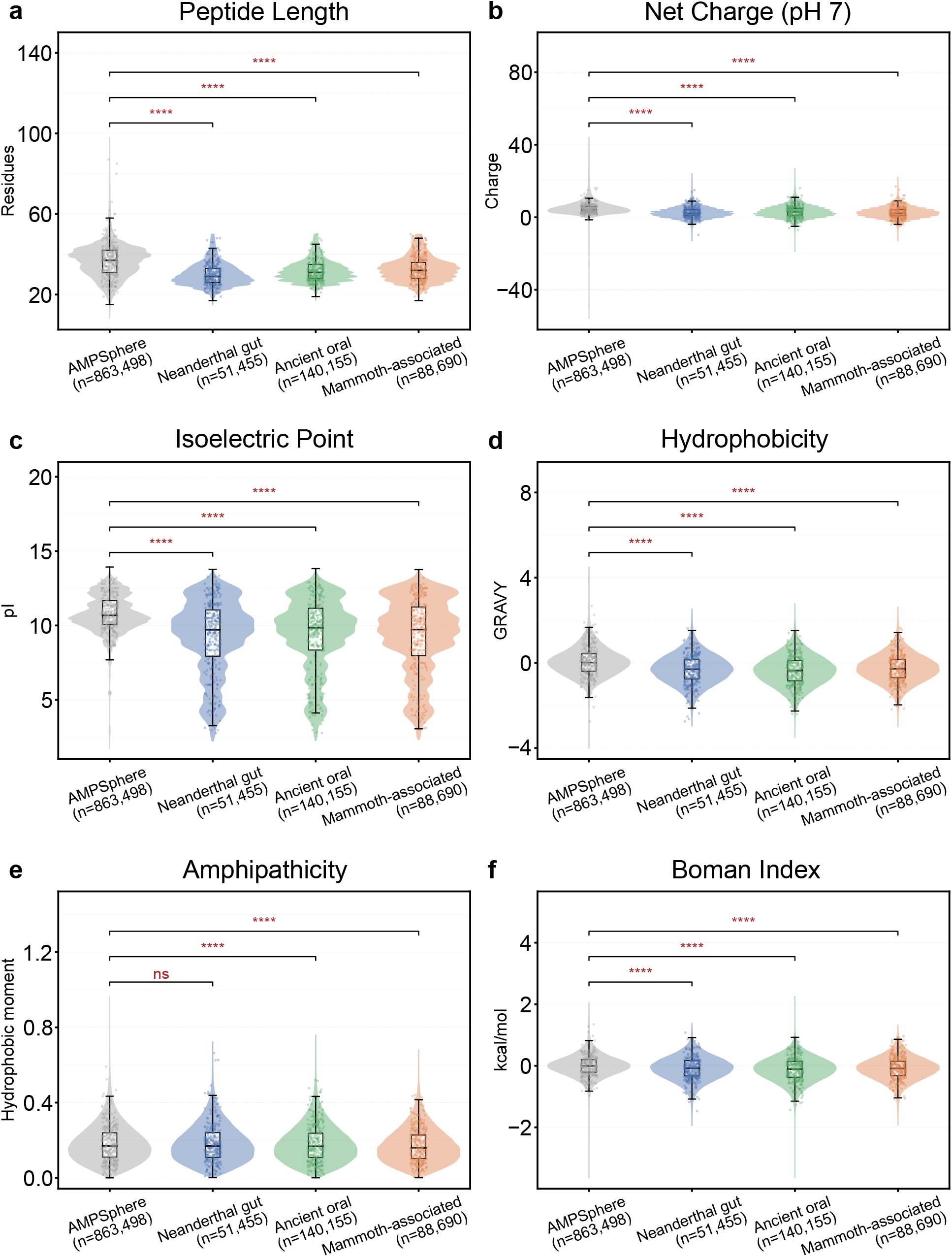
Physicochemical comparison of all ancient predicted AMPs with the modern AMPSphere catalogue. Six physicochemical properties are compared across four groups: the full AMPSphere catalogue of AMPs predicted from modern global microbiomes (n = 863,498) and all peptides predicted as antimicrobial by all three methods (APIN, EBiLSTM, and TabICL) from each of the three ancient datasets, the Neanderthal gut (n = 51,455), the ancient oral (dental calculus) microbiome (n = 140,155), and the mammoth-associated microbiome (n = 88,690). This comparison includes every peptide classified as antimicrobial at the standard 0.5 probability threshold. Panels show (a) peptide length, (b) net charge at pH 7, (c) isoelectric point, (d) hydrophobicity (GRAVY), (e) amphipathicity (Eisenberg hydrophobic moment), and (f) Boman index. For each group a violin shows the distribution, with an inner box plot indicating the median and interquartile range and individual peptides overlaid as points. Brackets indicate two-sided Mann-Whitney U tests comparing AMPSphere with each ancient dataset; given the large sample sizes, statistical significance reflects even small differences in distribution, so comparisons should be read through the location and spread of each distribution rather than through significance alone. Significance is denoted as: *, *p <* 0.05; **, *p <* 0.01; ***, *p <* 0.001; ****, *p <* 0.0001; ns, not significant.

We further examined the number of consensus candidates across increasingly stringent probability thresholds to guide high-confidence candidate selection. The candidate count fell by roughly an order of magnitude at each successive threshold in all three datasets. For the Neanderthal gut dataset, for example, it decreased from 25,866 at 0.9 to 8,326 at 0.99, 1,469 at 0.999, and 111 at 0.9999, and no candidates in any dataset passed the 0.99999 threshold. Based on this consistent pattern, we selected 0.9999 as a stringent cutoff, yielding 111 Neanderthal gut, 76 ancient oral, and 50 mammoth-associated candidates, representing 0.09%, 0.02%, and 0.02% of the respective non-redundant peptide sets. The high confidence candidates exhibited physicochemical properties consistent with known AMPs (**Figure 4**). Neanderthal gut, ancient oral, and mammoth-associated candidates had mean lengths of 23.0, 25.8, and 23.9 residues; mean net charges of +3.60, +4.42, and +3.82; mean isoelectric points of 10.50, 10.82, and 10.46; mean GRAVY values of -0.252, -0.131, and -0.209; and mean hydrophobic moments of 0.218, 0.215, and 0.219, respectively (Supplementary Table 2). These profiles indicate short, positively charged, and amphipathic peptides, consistent with membrane-active AMPs. Most candidates had a net charge of at least +2: 87/111 Neanderthal gut, 65/76 ancient oral, and 42/50 mammoth-associated peptides. Although not independent of the physicochemical information captured by protein language models, these characteristics provide a plausibility check for the predicted candidates.

**Figure 4.**
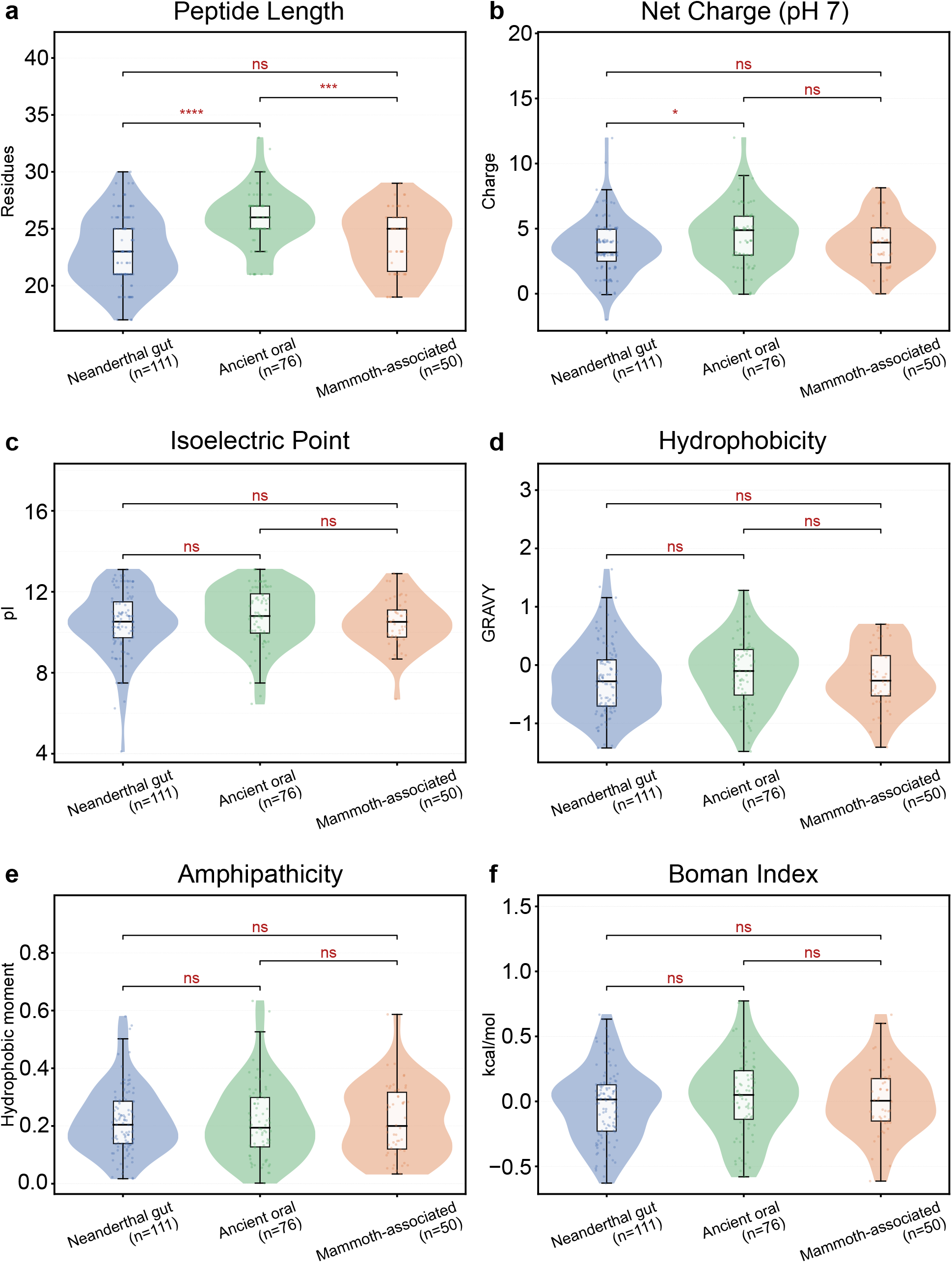
Physicochemical comparison of the high-confidence candidate antimicrobial peptides across the three ancient sources. Six physicochemical properties are compared among the high-confidence candidate peptides nominated from the three ancient datasets, the Neanderthal gut (n = 111), the ancient oral (dental calculus) microbiome (n = 76), and the mammoth-associated microbiome (n = 50). High-confidence candidates are the peptides assigned a probability of at least 0.9999 by all three methods (APIN, EBiLSTM, and TabICL). Panels show (a) peptide length, (b) net charge at pH 7, (c) isoelectric point, (d) hydrophobicity (GRAVY), (e) amphipathicity (Eisenberg hydrophobic moment), and (f) Boman index. For each group a violin shows the distribution, with an inner box plot indicating the median and interquartile range and individual peptides overlaid as points. Brackets indicate two-sided Mann-Whitney U tests between pairs of ancient sources. Significance is denoted as: *, *p <* 0.05; **, *p <* 0.01; ***, *p <* 0.001; ****, *p <* 0.0001; ns, not significant.

### AMP discovery in ancient microbiomes using reinforcement learning models without modern training bias

Supervised AMP prediction relies on models trained using modern AMP datasets, potentially limiting their ability to identify ancient peptides with distinct sequences and physicochemical properties. Therefore, we explore reinforcement learning as a complementary, reward-guided approach that does not rely on modern AMP labels. We compared candidates selected by PPO and GFlowNet under two reward designs, S1 based on amino-acid composition entropy and S2 on a physicochemical fitness function, yielding four candidate sets (PPO-S1, GFlowNet-S1, PPO-S2, GFlowNet-S2), against those identified by supervised models. Across the three datasets, S1 candidates were substantially longer and had lower net charges and isoelectric points than supervised predictions, whereas S2 candidates showed higher net charges and isoelectric points (**Figure 5**). Hydrophobicity differences were less consistent across datasets. Overall, reward design had a greater influence on candidate physicochemical profiles than the choice of search algorithm, supporting reward-guided selection as a complementary approach to exploring ancient peptide space.

**Figure 5.**
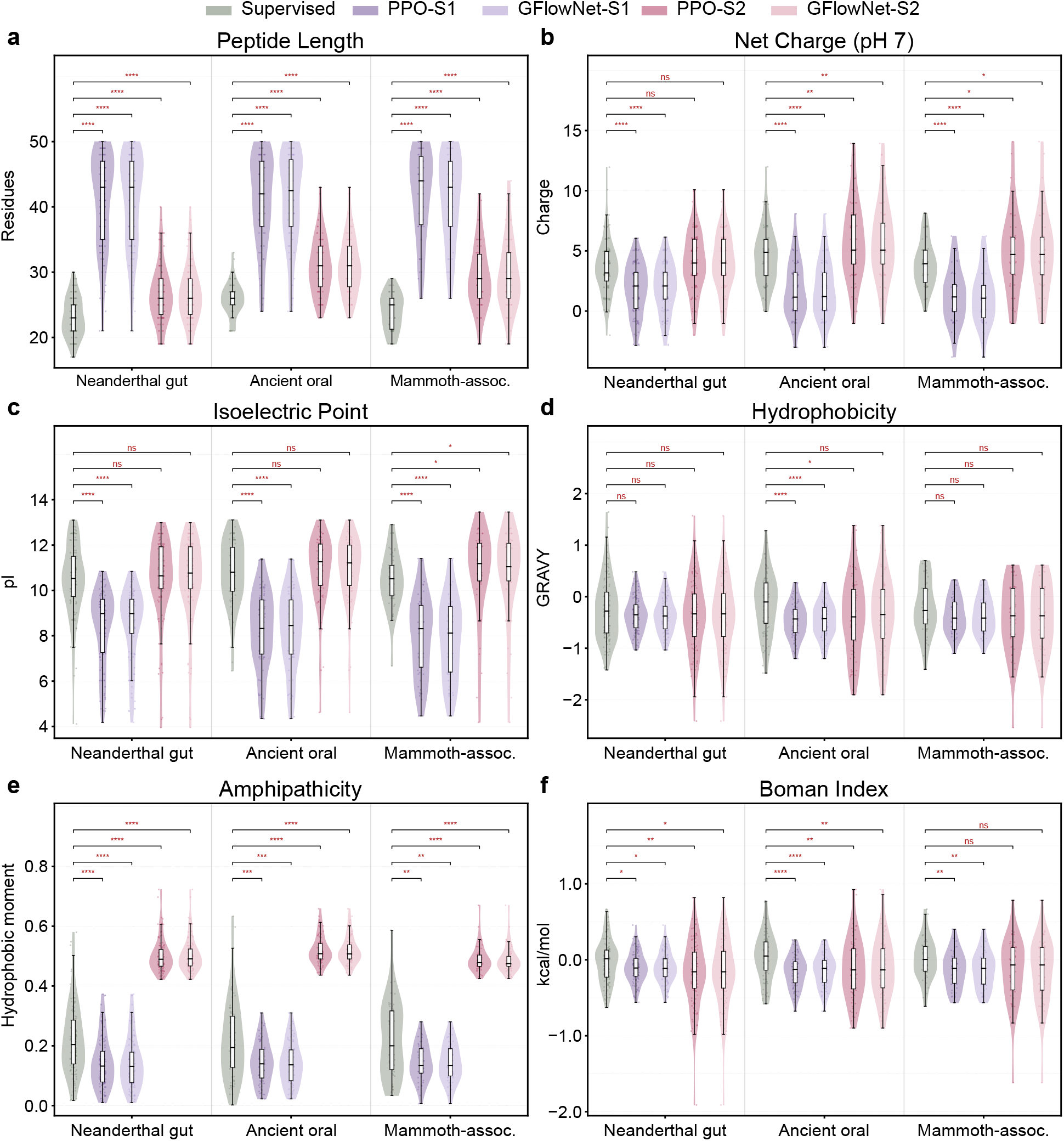
Physicochemical comparison of supervised and reinforcement-learning-selected candidate peptides. For each ancient dataset, the supervised high-confidence candidates are compared with peptides selected by PPO and GFlowNet under two reward designs: composition entropy (S1) and a physicochemical fitness function ^61^ (S2). Supervised candidates are the peptides assigned a probability of at least 0.9999 by all three methods (APIN, EBiLSTM, and TabICL). Each reinforcement-learning group is represented by a single random seed, matching the supervised candidate count; the three seeds were statistically interchangeable (Methods). PPO and GFlowNet are shown separately. All groups contain 111, 76, and 50 peptides for the Neanderthal gut, ancient oral, and mammoth-associated datasets, respectively. Panels show (a) peptide length, (b) net charge at pH 7, (c) isoelectric point, (d) hydrophobicity (GRAVY), (e) amphipathicity (Eisenberg hydrophobic moment), and (f) Boman index. For each group a violin shows the distribution, with an inner box plot indicating the median and interquartile range and individual peptides overlaid as points. Brackets indicate two-sided Mann-Whitney U tests between the supervised set and each reinforcement-learning group. Significance is denoted as: *, *p <* 0.05; **, *p <* 0.01; ***, *p <* 0.001; ****, *p <* 0.0001; ns, not significant.

## Discussion

This study provides a unified benchmark of machine learning methods for AMP prediction and applies the benchmarked pipeline to ancient microbiomes. The most consistent finding was that feature representation influenced performance more than model architecture: protein language model embeddings, particularly ESM-2 and ProtBERT^11, 27^, outperformed handcrafted descriptors across methods and datasets, and combining them with traditional features yielded only marginal gains. This indicates that language models already capture much of the sequence information relevant to antimicrobial activity, so effective prediction can be achieved without extensive feature engineering. The stability of method rankings across datasets of differing composition further suggests that these conclusions are not driven by dataset-specific characteristics. Practically, this lowers the barrier to AMP screening: a single pretrained embedding paired with a lightweight classifier can match more elaborate pipelines, making accurate prediction accessible without specialized feature design or large computational budgets.

Given strong sequence representations, architectural complexity provided limited additional benefit. The multi-scale convolutional model APIN^9^ performed best, consistent with antimicrobial activity being driven by motifs at several length scales^4^, yet simpler or training-free methods, including Random Forest^29^, a meta-tuned MLP^32^, and an in-context tabular transformer^33^, remained competitive. The graph network supplied with predicted residue contacts^30^ did not outperform sequence-only models, suggesting that explicit structural modeling offers little advantage for short peptides, which lack stable tertiary structure and whose relevant structural propensities may already be encoded in the embeddings^40^.

Applying this pipeline to three ancient metagenomes indicates that ancient microbiomes are an untapped source of candidate AMPs. The nominated peptides occupy the physicochemical space of known AMPs yet are largely distinct in sequence from modern references, with the degree of overlap depending strongly on body site: ancient gut and mammoth-associated sORFs were almost entirely novel, whereas most ancient oral sORFs had modern counterparts. This contrast is itself informative. The high oral overlap is consistent with the exceptional preservation of oral communities in dental calculus, which mineralizes around resident microbes, whereas the near-complete novelty of the gut and mammoth-associated sets points to sequence space not represented in current catalogs. Sequences that diverge from modern catalogs are of particular interest because novel scaffolds may act through mechanisms to which contemporary pathogens have not been exposed, though this potential remains to be tested. Ancient microbiomes may therefore harbor peptides with properties or mechanisms distinct from those already characterized, and reinforcement learning offers a complementary route to such candidates by searching ancient sequence space without modern AMP labels. These physicochemical similarities support the plausibility of the predictions but do not establish antimicrobial activity, which awaits experimental validation.

Several limitations qualify these findings. The benchmark addressed binary classification on eight datasets with some shared source databases and did not model computational cost or full structural information. The ancient candidates are computational predictions whose scores are best read as relative rankings rather than calibrated probabilities, and because supervised models are trained on modern AMPs, both the supervised and reinforcement-learning approaches may miss peptides outside their respective criteria. Finally, ancient metagenomes carry inherent risks of DNA degradation and contamination^25^; while the high oral overlap is consistent with good preservation in dental calculus, the low gut overlap could reflect genuine novelty or, in part, degradation or contamination, and the mammoth comparison is further limited because its samples span multiple anatomical sources whereas the modern reference is elephant gut.

These limitations point to natural extensions, including richer structural inputs, multi-label prediction of specific activities, domain-adapted language models, and few-shot in-context prediction for new peptide families^33^. Coupling the benchmarked predictors with generative design^41^, together with experimental validation of nominated candidates, would begin to translate these computational advances into functional antimicrobials. In summary, this work establishes a reproducible benchmark showing that protein language model embeddings with strong classifiers provide an accurate and generalizable basis for AMP prediction, and positions ancient microbiomes as a reservoir of candidate antimicrobials that predate modern antibiotics.

## Methods

We designed a systematic benchmark to evaluate how feature representations and prediction methods affect antimicrobial peptide identification. Each peptide is encoded as a set of per-residue features that combine traditional sequence descriptors with protein language model embeddings, and these representations are organized into distinct configurations that allow their individual and combined contributions to be assessed. The encoded peptides are then classified by six machine learning methods spanning tabular, convolutional, recurrent, and graph-based paradigms. The feature representations and prediction methods that constitute the benchmark are described below, along with the benchmark datasets and cross validation protocol used for evaluation, and the pipeline by which the best performing configuration was applied to three ancient metagenomes, a Neanderthal gut microbiome, an ancient oral microbiome, and a mammoth-associated microbiome recovered from skeletal and dental remains, to nominate and characterize novel AMP candidates.

### Feature representations

We extracted a comprehensive set of 258-dimensional features for each peptide sequence, combining traditional sequence descriptors with modern protein language model embeddings. These features are organized into six categories, enabling systematic ablation studies across different feature combinations.

### One-hot encoding (20 dimensions)

Each amino acid in the sequence is represented as a 20-dimensional binary vector, where each position corresponds to one of the 20 standard amino acids. For a given residue, the corresponding position is set to 1, and all other positions are set to 0. This captures the compositional information at each position without assuming any prior knowledge of amino acid properties.

### Sinusoidal Position Encoding (20 dimensions)

To capture positional information within the sequence, we employ sinusoidal position encoding^28^. For each position *pos* in the sequence, the encoding is computed as:

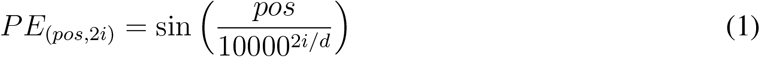

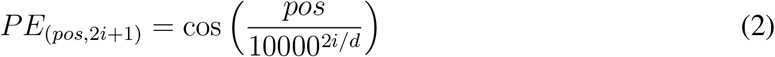

where *d = 20* is the encoding dimension and *i* ranges from 0 to 9. This lets the model distinguish residues at different positions and encodes their relative spacing.

### ESM-2 Embeddings (40 dimensions)

We utilize ESM-2 (Evolutionary Scale Modeling 2)^11^, a state-of-the-art protein language model pre-trained on millions of protein sequences. Specifically, we employ the esm2_t33_650M_UR50D model containing 650 million parameters. For each residue, we extract the 1280-dimensional embedding from the final layer and reduce it to 40 dimensions by Principal Component Analysis (PCA), which keeps the most informative components at a lower computational cost. The contact prediction capability of ESM-2 is also leveraged to generate predicted contact maps for graph-based methods.

### ProtBERT Embeddings (128 dimensions)

ProtBERT^27^ is a BERT-based protein language model pre-trained on UniRef100. We use the ’Rostlab/prot_bert’ model from Hugging Face, which produces 1024-dimensional embeddings. Following extraction, we apply a linear projection layer to reduce the dimensionality to 128. The input sequences are tokenized by adding spaces between amino acids, and rare amino acids (U, Z, O, B) are replaced with X. Embeddings are extracted from the last hidden state, excluding the [CLS] and [SEP] tokens.

### Amino Acid Composition (20 dimensions)

Amino Acid Composition (AAC) represents the frequency of each of the 20 standard amino acids within a sequence. For each sequence, we calculate the normalized count of each amino acid, resulting in a 20-dimensional feature vector. This global compositional signature is then replicated for each residue position, providing consistent sequence level information at the residue level for compatibility with residue level models.

### Physicochemical Properties (30 dimensions)

We encode six fundamental physicochemical properties of amino acids: hydrophobicity (Kyte-Doolittle scale), charge, polarity, aromaticity, size (molecular weight), and secondary structure propensity (alpha helix tendency). For each property, we compute five statistical measures—mean, standard deviation, minimum, maximum, and sum—yielding a 30-dimensional feature vector. To capture local context, we calculate position specific features using a sliding window of size 7, combining global (60%) and local (40%) statistics.

### Contact Maps

For graph-based methods, we derive predicted contact maps from the ESM-2 contact prediction head. Contact maps represent pairwise residue interactions as a binary adjacency matrix, where an edge exists between residues if their predicted *C_β_ − C_β_* distance is below 8Å. These contact maps are used as edge information for the Graph Convolutional Network (GCN), while other methods operate solely on the node (residue) features.

### Feature configurations

From these feature types we defined twelve configurations for evaluation. Rather than exhaustively testing all 63 possible non-empty combinations of the six feature types, which would be combinatorially prohibitive and largely redundant, we selected twelve representative configurations: the six individual feature types, three combinations pairing related descriptors (one-hot with position encoding, amino acid composition with physicochemical properties, and the two protein language model embeddings), and three combinations that progressively integrate handcrafted and language model features. The full list of configurations is given in Table 1.

### Prediction methods

We benchmark six machine learning methods spanning traditional models, deep learning architectures, and modern transformer based approaches. These methods represent distinct paradigms in sequence classification: tree based ensembles that operate on engineered features, graph neural networks that leverage predicted structural information, convolutional networks that capture local sequence motifs, recurrent networks that model long range dependencies, and tabular foundation models that exploit in-context learning. All methods are evaluated under a unified experimental framework with consistent data splits, training protocols, and evaluation metrics to ensure fair comparison. Full architectural specifications, equations, and hyperparameters for each method are provided in Supplementary Note 1.

### Random Forest

Random Forest (RF) is an ensemble method that builds many decision trees and classifies by majority vote^29^. We use it as a strong non deep learning baseline (scikit-learn implementation). Because RF requires fixed length inputs while peptides vary in length, we aggregate the residue level features to a sequence-level vector by mean pooling, the most widely adopted and empirically strongest compression for variable length protein language model outputs^42^. The classifier uses 300 trees with unlimited depth and square root feature sampling at each split. RF operates on these mean pooled sequence level representations rather than residue level matrices.

### Graph Convolutional Network (GCN)

GCN operates on graph-structured data, making it suited to modeling residue interactions^30^. We build a graph per peptide in which nodes are residues carrying their feature vectors and edges are derived from ESM-2 predicted contact maps, so that, unlike the other methods, GCN explicitly uses predicted structural information. The architecture stacks three graph convolutional layers (hidden dimension 128), each with batch normalization, ReLU, and dropout, followed by global mean pooling to a graph level representation and a two-layer MLP classifier. Each layer performs neighborhood aggregation over the contact derived adjacency (propagation rule and full settings in Supplementary Note 1).

### APIN (CNN-based)

APIN (Antimicrobial Peptide Identification using a multi-scale convolutional Network) uses parallel convolutional filters of different sizes to capture sequence motifs of varying length^9^, since functional peptide motifs range from short cationic patches to longer amphipathic stretches. The network applies 14 parallel 1D convolutions with filter sizes from 2 to 28 (64 filters each), each followed by batch normalization, ReLU, and max pooling; the resulting multi-scale features are combined by global max pooling across scales into a 64-dimensional representation, which a two-layer classifier maps to the AMP/non-AMP decision. This design lets the model detect short and long motifs simultaneously.

### EBiLSTM (Bi-LSTM)

EBiLSTM is an enhanced bidirectional LSTM for AMP prediction^31^. It processes residue level feature sequences through three stacked bidirectional LSTM layers with decreasing hidden sizes (128, 64, 32), each with dropout, so that sequence context is read from both the N- and C-terminal directions and summarized hierarchically. The final representation is fed into a two-layer classifier for the binary decision. Unlike the tabular methods, EBiLSTM operates directly on residue level feature matrices (padded or truncated to 200 residues), using packed sequences so padded positions are ignored, and is trained with Adam and early stopping on validation AUC. As no source code was available, we reimplemented it in PyTorch from the published specification.

### TabICL (Transformer)

TabICL is a transformer based tabular foundation model that performs in-context learning: rather than updating parameters per task, it conditions on labeled examples in its context window to classify query samples^33^. It is pretrained on a large, diverse collection of synthetic classification tasks, which lets it transfer to new tasks without gradient based training. Like RF, it requires fixed length inputs and therefore operates on mean pooled sequence level representations. To improve robustness and calibration we use an ensemble of 32 members with different context samplings, averaging their predicted probabilities. This in-context approach makes TabICL a training free counterpart to the other classifiers.

### RealMLP

RealMLP is a multilayer perceptron optimized for tabular data, with architecture, preprocessing, regularization, and training defaults meta tuned across 118 diverse datasets for strong out-of-the-box performance without per-task tuning^32^. Like RF and TabICL, it uses mean pooled sequence level representations. The network has three hidden layers of 256 units; its meta tuned defaults (learning rate, weight decay, batch size, epochs, initialization, and feature normalization) are detailed in Supplementary Note 1. RealMLP serves as a strong, modern feedforward baseline that contrasts with both the tree ensemble and the sequence aware deep models.

### Training datasets

To objectively evaluate the performance of the prediction methods, we utilize eight independent test datasets commonly used for AMP prediction, including XUAMP^6^, APD3^6, 43^, DRAMP^44, 45^, LAMP^46, 47^, CAMP^13, 48^, dbAMP^49^, YADAMP^50^, and DBAASP^51^. These datasets were originally compiled and curated by Xu et al.^6^ and Yan et al.^52^, with each dataset consisting of balanced positive (AMP) and negative (non-AMP) samples to enable fair comparison across different prediction methods.

Specifically, XUAMP is the largest independent test dataset, containing 8,572 positive and 8,572 negative samples. This dataset was constructed by Xu et al.^6^ based on multiple AMP databases including DRAMP, LAMP, and YADAMP, with redundancy removal to ensure sequence diversity. APD3 is derived from the Antimicrobial Peptide Database version 3^43^, one of the earliest and most widely used AMP databases, containing 6,745 positive and 6,682 negative samples. DRAMP is sourced from the Data Repository of Antimicrobial Peptides^44, 45^, comprising 7,369 positive and 6,958 negative samples. DRAMP provides comprehensive annotations including sequence, activity, and structural information. LAMP is from the Linking Antimicrobial Peptides database^46, 47^, containing 6,868 positive and 6,878 negative samples. LAMP focuses on linking AMPs with their functional and structural properties. CAMP is derived from the Collection of Anti-Microbial Peptides^13, 48^, with 6,707 positive and 6,597 negative samples. This is a highly curated dataset. dbAMP is from the database of Antimicrobial Peptides^49^, containing 6,719 positive and 6,688 negative samples. dbAMP integrates transcriptome and proteome data for comprehensive AMP analysis. YADAMP (Yet Another Database of Antimicrobial Peptides)^50^ comprises 6,755 positive and 6,630 negative samples, providing detailed physicochemical property annotations. DBAASP is from the Database of Antimicrobial Activity and Structure of Peptides^51^, containing 7,214 positive and 7,214 negative samples. Following Yan et al.^52^, this dataset was constructed with positive samples sharing less than 90% sequence similarity and negative samples sharing less than 40% similarity.

The statistical information of all eight datasets is summarized in Table 3. While most positive samples across datasets are unique, there exists some overlap among certain datasets, as recent datasets were often constructed based on earlier ones. For example, XUAMP was constructed by aggregating sequences from DRAMP, LAMP, and YADAMP^6^. XUAMP and DRAMP represent the two largest datasets with similar sequence length distributions between positive and negative samples, making them particularly suitable as primary evaluation benchmarks. The remaining six datasets exhibit varying sequence length distributions, providing a comprehensive evaluation across diverse data characteristics.

**Table 3:** Summary statistics of the eight benchmark datasets.

| **Dataset** | **Positive** | **Negative** | **Total** |
|---|---|---|---|
| XUAMP | 8,572 | 8,572 | 17,144 |
| APD3 | 6,745 | 6,682 | 13,427 |
| DRAMP | 7,369 | 6,958 | 14,327 |
| LAMP | 6,868 | 6,878 | 13,746 |
| CAMP | 6,707 | 6,597 | 13,304 |
| dbAMP | 6,719 | 6,688 | 13,407 |
| YADAMP | 6,755 | 6,630 | 13,385 |
| DBAASP | 7,214 | 7,214 | 14,428 |
| **Total** | 56,949 | 56,219 | 113,168 |

### Model evaluation setup

All methods are evaluated using 10-fold cross validation with stratified sampling to maintain class balance. For each fold, 80% of the data is used for training and 20% for testing. We report the mean and standard deviation of all metrics across the 10 folds to assess both performance and stability. We evaluate model performance using six standard metrics: Area Under the ROC Curve (AUC), Accuracy (ACC), F1-Score, Matthews Correlation Coefficient (MCC), Sensitivity (Sn, recall), and Specificity (Sp). All experiments are implemented in Python using PyTorch for deep learning models and scikit-learn for traditional machine learning methods. Graph neural network operations utilize PyTorch Geometric. Protein language model embeddings are extracted using the Hugging Face Transformers library for ProtBERT and the ESM library for ESM-2. All deep learning methods follow a consistent two-stage training protocol. In Stage 1 (Pretraining), models are trained on the pretrain split with validation on a held out portion. In Stage 2 (Fine tuning), models are trained on the finetune split. The final model is selected based on the higher validation AUC between the two stages. This protocol ensures fair comparison across methods while allowing models to benefit from the full training data. To assess whether performance differences among the six methods were statistically significant, we applied a Friedman test across the eight datasets to the per-dataset mean AUCs at the best performing feature configuration, followed by a Nemenyi post-hoc test at *α* = 0.05. Tests were computed in Python using SciPy for the Friedman test and the average rank and critical difference calculations for the Nemenyi procedure.

### Ancient metagenomic datasets

We applied the benchmarked pipeline to three ancient metagenomic datasets spanning human and mammoth hosts and three distinct sample types. The Neanderthal gut dataset derives from Middle Paleolithic fecal sediments from the El Salt site in Spain, dated to approximately 50,000 years ago^34^. The ancient oral dataset derives from archaeological dental calculus from human skeletal remains spanning the Neolithic to the Early Middle Ages^35^. The mammoth-associated dataset derives from ancient microbial DNA recovered from mammoth skeletal and dental remains spanning more than one million years^36^. Raw shotgun metagenomic sequencing data for the three cohorts were obtained from the European Nucleotide Archive (ENA) under accession numbers PRJEB41665, PRJEB43389, and PRJEB78615.

Metagenomic sequences were processed following a uniform pipeline. Raw reads for each sample were quality controlled with MetaWRAP (v1.3.2)^53^, including adapter and quality trimming and removal of host-derived reads. Reads from the two ancient human microbiome datasets were mapped against the CHM13^54^ human reference genome for host-read removal, whereas reads from the mammoth-associated dataset were mapped against the Elephas maximus genome and CHM13 to remove host-derived sequences. The resulting clean reads were assembled with the metaWRAP-Assembly module using metaSPAdes (v3.13.0). Small open reading frames (sORFs, 10 to 50 residues) were then predicted using the ’getorf’ function of the EMBOSS software package (version 6.6.0)^55^. To keep peptides that recur across samples rather than one-off or spurious calls, we ranked the candidates in each dataset by their prevalence across that cohort’s samples and retained only those detected in at least 10% of samples, or in at least 5 independent samples, whichever criterion was less stringent.

### sORFs clustering

Following prevalence filtering, the number of retained sORFs remained large and computationally challenging for downstream AMP prediction. To reduce sequence redundancy and computational burden, we clustered the prevalence filtered peptides using CD-HIT^56^ at a 50% sequence identity threshold (word size 3, minimum sequence length 5), retaining one representative sequence per cluster. A 50% threshold was selected after comparing 90%, 70%, and 50% identity cutoffs: higher thresholds collapsed the sets only marginally, reflecting the high sequence diversity of these ancient peptide pools, whereas 50% grouped sequences at the family level while remaining within the range for which CD-HIT is recommended for protein clustering. Clustering was applied after, rather than before, any length based filtering, because CD-HIT selects the longest member of each cluster as its representative and clustering-first would therefore systematically deplete the shortest peptides. This clustering yielded 118,583, 387,477, and 287,622 non-redundant candidate peptides for the Neanderthal gut, ancient oral, and mammoth-associated datasets, respectively.

### Modern reference sORF set and sequence overlap analysis

To assess whether the ancient sORFs are distinct in sequence from their modern counterparts, we constructed a body-site-matched modern reference for each ancient dataset. The Neanderthal gut set was compared against the Human Reference Gut Microbiome v2 (HRGM2), comprising 4,824 representative genomes^37^. The ancient oral set against the Human Reference Oral Microbiome (HROM), comprising 5,113 representative genomes^38^. And the mammoth-associated set against 734 elephant gut metagenome-assembled genomes obtained from the gcMeta repository^39^, the closest available modern analogue. Each reference was processed identically: small open reading frames were predicted with the `getorf` program of EMBOSS (version 6.6.0)^55^ using the same settings as for the ancient datasets (`-find 0`, bacterial translation table 11, 30 to 150 nucleotides, corresponding to 10 to 50 residues), then clustered per genome with CD-HIT^56^ at 50% identity and pooled into a non-redundant modern set (141,253,669, 108,621,857, and 21,728,079 sORFs for the gut, oral, and elephant references, respectively). Each non-redundant ancient sORF set was then compared against its matched modern reference with `cd-hit-2d` at 90% identity, and we report the fraction of ancient sORFs with at least one modern match. Full counts are given in Supplementary Note 2.

### Training set construction and prediction on ancient peptides

Because the ancient peptides are unlabeled, we trained a single model on all available labeled data rather than performing cross validation. We pooled the positive and negative sequences from all five splits of the eight benchmark datasets, giving 113,168 records, and removed duplicates at the sequence level. The eight datasets overlap substantially, since several were derived from shared source databases, and deduplication reduced the pool to 15,475 unique sequences. A further 36 sequences carried conflicting labels across datasets, appearing as AMP in one and non-AMP in another, and were discarded. The final training set comprised 15,439 peptides (8,097 AMP and 7,342 non-AMP).

For the ancient application we combined the two central findings of our benchmark: the three best performing methods and the single most informative feature representation. Method ranking was established at the best performing configuration, where APIN^9^, TabICL^33^, and EBiLSTM^31^ were the three strongest models. As the feature representation we used ESM-2 embeddings alone (Configuration 3), which were the most informative single feature in our benchmark (average AUC 0.9718). Restricting the ancient application to this single embedding, rather than the full multi-feature combination, substantially reduces the computational cost of scoring hundreds of thousands of candidate peptides while retaining nearly all of the predictive performance, and keeps the feature space identical across the benchmark and ancient datasets. Features were extracted exactly as described above, using the same PCA projection fitted on the benchmark data so that all sequences occupy a common 40-dimensional space. These three methods were each trained on the combined training set with a stratified 10% held out partition used solely for early stopping, then applied to each of the three ancient datasets to obtain per-peptide AMP probabilities. A peptide was retained as a high confidence candidate only if all three methods assigned it a probability of at least 0.9999. This threshold was chosen as a deliberately conservative operating point. Across all three datasets the number of consensus candidates decreased by approximately an order of magnitude per threshold decade and reached zero at *≥*0.99999, indicating that 0.9999 isolates the extreme high confidence tail of the score distribution.

### Physicochemical characterization

For each high confidence candidate we computed a panel of physicochemical descriptors using a custom Python implementation based on standard published scales, rather than default parameters of any single software package, so that every value is fully specified and reproducible. Net charge was calculated at pH 7.0 by summing the protonation states of all ionizable groups (the N- and C-termini and the Asp, Glu, Cys, Tyr, His, Lys, and Arg side chains) according to the Henderson–Hasselbalch equation using standard side-chain p*K_a_*values, and the isoelectric point was obtained as the pH at which the net charge is zero, located by bisection on the same charge model. Mean hydrophobicity was computed as the average of the Kyte–Doolittle hydropathy scale (GRAVY)^57^. Amphipathicity was quantified by the mean hydrophobic moment^58^, using the Eisenberg consensus hydrophobicity scale and an assumed *α*-helical periodicity of 100*^◦^* per residue. We additionally computed the aliphatic index^59^, the Boman (protein binding potential) index^60^, molecular weight, charge density (net charge per residue), aromaticity (the fraction of Phe, Trp, and Tyr), and the fractions of cationic (Lys, Arg) and hydrophobic (Ala, Cys, Phe, Ile, Leu, Met, Val, Trp, Tyr) residues. None of these descriptors were provided to the prediction models, which operated on ESM-2 embeddings alone; they therefore constitute an independent characterization of the selected peptides.

### Reinforcement-learning candidate selection

We applied proximal policy optimization (PPO) and GFlowNet to the same ancient peptide pools used in the supervised analysis. Sequence descriptors were standardized, reduced to 64 dimensions by PCA, and L2-normalized; each trajectory selected 12 distinct peptides. Two reward designs were used: S1, based on normalized amino-acid composition entropy, and S2, based on a physicochemical fitness function for amphipathic *α*-helical peptides^61^. The subset score was

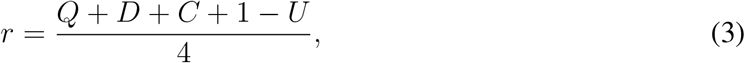

where *Q* is sequence quality, *D* is feature dissimilarity, *C* is the number of represented clusters divided by subset size, and *U* penalizes high feature similarity. After subset evaluation, the unique sequences encountered during the search were ranked by their quality score *q*, retaining the top 111, 76, and 50 for the Neanderthal gut, ancient oral, and mammoth-associated datasets, respectively, to match the supervised candidate counts. AMP-predictor scores and pretrained protein embeddings were not used during the search. Because candidate search is stochastic, each method and reward was run with three random seeds (17, 29, and 43). The seeds produced statistically indistinguishable physicochemical distributions, 72 Kruskal-Wallis tests across the two methods, two rewards, three datasets, and six properties, with none significant after Benjamini-Hochberg correction, so a single seed was used to represent each group in the main comparison.

## Data Availability

The metagenomic sequencing data in this study can be downloaded via ENA (PRJEB41665, PRJEB43389, and PRJEB78615). Training data of our machine learning models are derived from publicly available antimicrobial peptide databases: dbAMP (http://awi.cuhk.edu.cn/dbAMP), DRAMP (http://dramp.cpu-bioinfor.org/), DBAASP (https://dbaasp.org/), APD3 (https://aps.unmc.edu/), and CAMP (https://camp3.bicnirrh.res.in/), together with the LAMP^46^, YADAMP^50^, and XUAMP^6^ datasets. Negative samples were obtained from UniProt (https://www.uniprot.org/).

## Code Availability

The code used in this study for model training, prediction, statistical analysis, and visualization is available in the GitHub repository (https://github.com/KelabatOSU/amp-benchmark_ancient_mining).

## Author Contribution

S.K. and C.C. conceived and designed the project. Q.Z. performed all the data analysis. Q.Z., S.K. and C.C. interpreted the results and prepared the manuscript. All authors approved the manuscript.

